# Activation of the noncanonical inflammasome-GSDMD pathway triggers pyroptosis in bone marrow and promotes periosteal bone formation

**DOI:** 10.1101/2025.10.20.683492

**Authors:** Wei Zou, Chun Wang, Yongjia Li, Wentong Jia, Steven L Teitelbaum, Gabriel Mbalaviele

## Abstract

Evidence indicating that inflammation is commonly associated with ectopic osteogenesis in certain autoimmune and infectious conditions challenges the dogma that inflammatory responses always suppress bone formation. In this study, we find that systemic administration of lipopolysaccharide (LPS) to mice causes not only inflammation in bone marrow, as expected, but also stimulates periosteal bone formation. This response can be reproduced *in vitro* as bone marrow supernatants from LPS-treated mice induce robust osteogenesis of skeletal stem cells (SSCs) compared to supernatants from PBS-treated counterparts. Periosteal bone accrual is partly dependent on periosteal leptin receptor-positive (LepR)^+^ SSCs but not bone marrow LepR^+^ or adiponectin (Adq)^+^ SSCs and correlates with pyroptosis within bone marrow. Consistent with the dependence of periosteal osteogenesis on pyroptosis, this response is slightly attenuated in *Nlrp3^-/-^* or *caspase-1^-/-^*mice but significantly inhibited in *caspase-11^-/-^*, *caspase-1^-/-^*;*caspase-11^-/-^*, or *Gsdmd^-/-^*mice. Our study reveals a novel role for pyroptosis in which lysed cells release intracellular contents that stimulate osteoprogenitors and promote osteogenic differentiation within the periosteal compartment.

## Introduction

Bone mass is decreased in many diseases, including osteoporosis, inflammatory arthritis and malignancies, and is associated with increased risks for the development of late fractures, and morbidities in the elderly populations^1–3^. In fact, it was reported that 20-50% of geriatric patients (≥65 years) with a hip fracture die within one year of fracture^4^. Although significant progress has been made in the treatment of low bone mass disorders, there remains an unmet medical need for the development of therapeutic agents that specifically improve the strength of cortical bone, which comprises most of skeletal bone mass (80%) at sites such as the hip and wrist, and provides approximately 65-90% of proximal femur strength^5–7^. A better understanding of nuances between these processes will lead to the development of safer and effective therapeutics for the treatment of disorders such as osteoporosis and delayed fracture healing.

Bone mass and quality are maintained throughout the lifespan by a tightly balanced process mediated by bone-resorbing osteoclasts (OCs) and bone-forming osteoblasts (OBs)^8^. While OCs arise from hematopoietic stem cells (HSCs), OBs originate from SSCs^9–15^. Within adult bone marrow, leptin receptor-expressing (LepR) SSCs are the major source of new OBs for bone maintenance^13^. These cells or their subsets also express adiponectin (Adq), Cxcl12, or Ebf3^11, 12, 14–16^. An imbalance whereby bone resorption outpaces bone formation underlies the pathogenesis of diseases of bone loss^17^. The current dogma posits that inflammation tilts this balance in favor of bone resorption and decreased bone formation^9, 18^. This view is supported by clinical data indicating that high levels of inflammatory factors such as interleukin (IL)-6 and tumor necrosis factor (TNF)-α are associated with a higher risk of bone fracture, and overwhelming preclinical evidence indicating that these cytokines stimulate bone resorption while inhibiting bone formation ^19–21^. Contrasting these findings are numerous studies showing that inflammation can promote osteogenesis. Indeed, several human infectious and non-infectious inflammatory conditions, such as ankylosing spondylitis^22, 23^, osteomyelitis^24,25^, and fracture healing^26^ are associated with cortical bone accrual. In support of this clinical evidence, experimental osteomyelitis in which *Staphylococcus aureus* is injected into femoral diaphysis is characterized by extensive cortical bone loss associated with robust reactive cortical bone formation surrounding the site of infection^27, 28^. Similarly, administration of TLR2 or TLR4 agonists into mouse calvaria induces bone formation through canonical Wnt signaling pathways^29^. In sum, although inflammation promotes bone resorption, it can give rise to bone formation, an outcome that is probably tissue context dependent. Coincidentally, inflammation predominantly induces reactive periosteal bone formation^23, 27, 28, 30^.

Emerging research implicates lytic forms of programed cell death, including pyroptosis, necroptosis, and PANoptosis in the propagation of inflammation^31, 32^. Pyroptosis is caused by cell stressors such as LPS, a component of the outer membrane of gram-negative bacteria and is mediated by a family of proteins called gasdermins (GSDMs) of which GSDMD is the most studied family member^33–36^. Cleavage of GSDMD by caspase-1, caspase-11, or neutrophil elastase generates GSDMD amino-terminal fragments, which form plasma membrane pores^33–36^. GSDMD conduits facilitate the secretion of IL-1β and IL-18 by live cells, but excessive pore formation can compromise the integrity of the plasma membrane and causes pyroptosis, leading to the release of not only IL-1β and IL-18, but also intracellular contents^35, 36^. Thus, GSDMD pores can exacerbate inflammation through pyroptosis-dependent mechanisms. We and others have previously reported that inflammasome-GSDMD pathways play an important role in bone development and maintenance in humans and rodents^37–41^. Consistent with its functional link to inflammasomes, LPS stimulates bone resorption by upregulating OC differentiation and activity in disease models^42^. LPS is found in systemic circulation in patients with increased gut permeability or decreased liver functions, and has been linked to multiple conditions, including skeletal diseases^43–47^. Thus, LPS, inflammasomes, and GSDMD are involved in bone metabolism. However, the extent to which pyroptosis per se, impacts bone pathophysiology is unknown.

In this study, we have identified a novel mechanism in which LPS activates the non-canonical inflammasome pathway, leading to GSDMD-mediated pyroptosis in bone marrow myeloid cells. This response ultimately induces periosteal LepR^+^ cells to differentiate into OBs and form bone.

## Results

### LPS stimulates periosteal bone formation, a response that is decoupled from bone resorption

We previously generated *Dtr^Adq^* mice bearing primate diphtheria toxin receptor (Dtr) downstream of a floxed-stop codon and expressing *Adq-Cre*. We found that elimination of Adq^+^ cells upon injection of diphtheria toxin (DT) caused systemic increased of bone mass^48^. To determine the role that inflammation plays in this response, we exposed DT-treated *Dtr^Adq^* mice to a low dose of *E*. *coli*-derived LPS for 10 days. While LPS administration decreased bone formation in bone marrow cavity of Adq^+^ cell-ablated mice, it dramatically and uniquely promotes periosteal bone formation (Supplementary Fig. 1A). To rule out the influence of DT in this response, we subjected wild-type (WT) mice to a similar LPS regimen. While LPS had no effect on the endosteal bone envelope, it induced massive periosteal bone formation (Fig. 1A, B). While LPS significantly enhanced cortical bone parameters, including thickness and area as well as periosteal perimeter, it did not affect bone marrow area, endocortical perimeter, and trabecular bone (Fig. 1A, B and Supplementary Fig. 1B, C). Periosteal osteogenesis also occurred in the tibia, humerus, and ilium (Supplementary Fig. 2). To ensure that induction of periosteal osteogenesis was not limited to *E*. *coli*-derived LPS, mice were also side-by-side inoculated with an identical dose of LPS (1 mg/kg) from *E. coli* or *Salmonella enterica*. *Salmonella*’s LPS increased cortical bone thickness without affecting trabecular bone mass in a similar manner to *E. coli*’s LPS (Supplementary Fig. 3A, B). The similarity of bone alterations induced by LPS from two different bacterial strains suggest that these responses are driven by LPS-intrinsic properties.

**Fig. 1.**
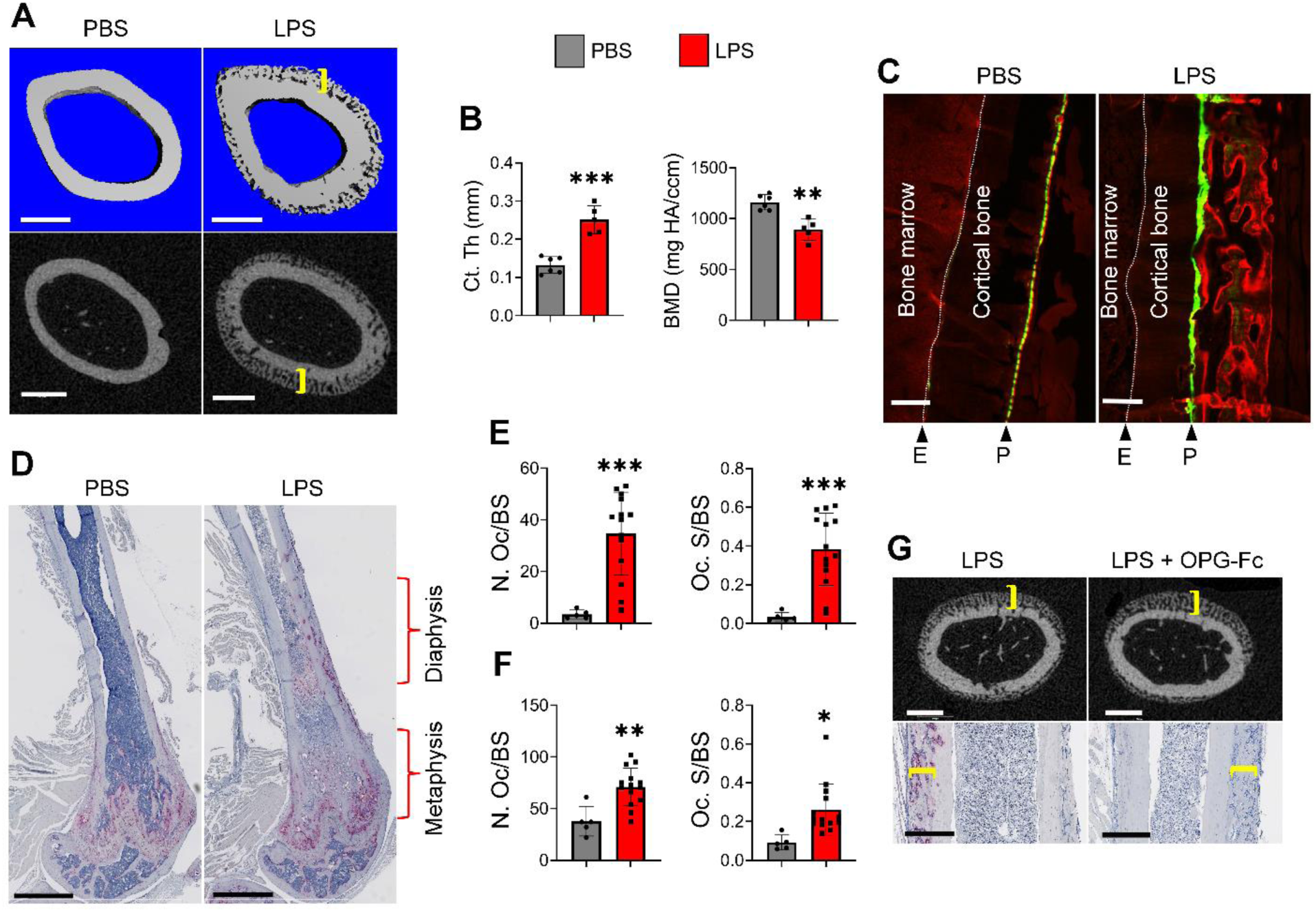
LPS stimulates periosteal bone formation, a response that is decoupled from bone resorption. **A**-**F**) Three months old WT mice were i.p. injected with PBS or 1 mg/kg LPS from *E. coli* on day 0, 4, and 8, mice were sacrificed on day 10. **A**) Representative μCT images of femoral diaphysis treated with PBS (n=6) and LPS (n=5). Scale bar: 500 µm. **B**) Cortical thickness (Ct.Th) and bone mineral density (BMD) analysis of **A**). **C**) WT mice were i.p. injected with PBS (n = 5) or LPS (n = 11) on day 0, 4, and 8, then with calcein and alizarin red on day 4 and 8, respectively. Representative images of calcein- and alizarin red-labelled femurs. Scale bar: 200 µm. E, endosteum; P, periosteum. **D**-**F**) Femoral sections were stained for TRAP activity. **D**) Representative images. Scale bar: 1 mm. **E**) Quantitative analysis of diaphyseal areas. **F**) Quantitative analysis of metaphyseal areas. N.Oc/BS, OC number/bone surface. Oc.S/BS, OC surface/bone surface. **G**) Three months old WT mice were i.p. injected with 1 mg/kg LPS with (n=10) or without OPG-Fc (5 mg/kg/mouse; n=10) on day 0, 4, and 8. Mice were sacrificed on day 10 and femurs were collected for µCT analysis (top; scale bar: 500 µm) and TRAP staining (bottom; scale bar: 400 µm). Brackets indicate the thickness of the newly formed woven bone, which was histologically identified by its characteristic porous appearance. *p<0.05, **p<0.01, ***p<0.001. Unpaired t test.

Focusing on *E*. *coli*-derived LPS, calcein and alizarin red double labelling experiments confirmed accrual periosteal but not endosteal bone formation in response to LPS treatment (Fig. 1C). We also unexpectedly found that LPS decreased cortical bone mineral density (Fig. 1B), presumably due to increased OC-mediated bone resorption. Consistent with this view, LPS enhanced OC number and surface not only in metaphyseal, but also in diaphyseal bone regions (Fig. 1D-F). Thus, LPS induced bone resorption throughout the femur, but reactive bone formation occurred mainly in the diaphyseal area. To determine whether periosteal bone formation and resorption induced by LPS were coupled, we injected WT mice with LPS in the presence or absence of osteoprotegerin-Fc (OPG-Fc) to block OC differentiation. While administration of OPG-Fc completely inhibited OC formation in both metaphyseal and diaphyseal bone regions, it did not affect periosteal osteogenesis (Fig. 1G and Supplementary Fig. 4). To determine whether the periosteal woven bone undergoes remodeling into mature bone, both PBS- and LPS-exposed samples were analyzed not only on day 10, but also on days 17 and 28. Periosteal bone thickness increased on days 10 and 17 following LPS exposure compared to PBS-treated samples, before declining slightly by day 28 (Fig. 2A, B). This increase correlated with the presence of OCs within the woven bone, which peaked on days 10 and 17 and remained at intermediate levels on day 28, compared to the PBS group. Although BMD declined on days 10 and 17, it returned to baseline by day 28. Interestingly, by day 28, LPS-treated mice exhibited clear evidence of bone trabeculation (Supplementary Fig. 5). Further biomechanical analyses revealed that key parameters, including stiffness, yield load, maximum load, and work to fracture, were all elevated in LPS-treated mice compared to PBS-treated controls on day 28 (Fig. 2C). Together, these findings suggest that periosteal bone formed in response to LPS undergoes remodeling and integrates with the existing bone, remains stable over time as the formation of new trabeculae is initiated. Ultimately, the overall bone structure is stronger relative to its pre-LPS state.

**Fig. 2.**
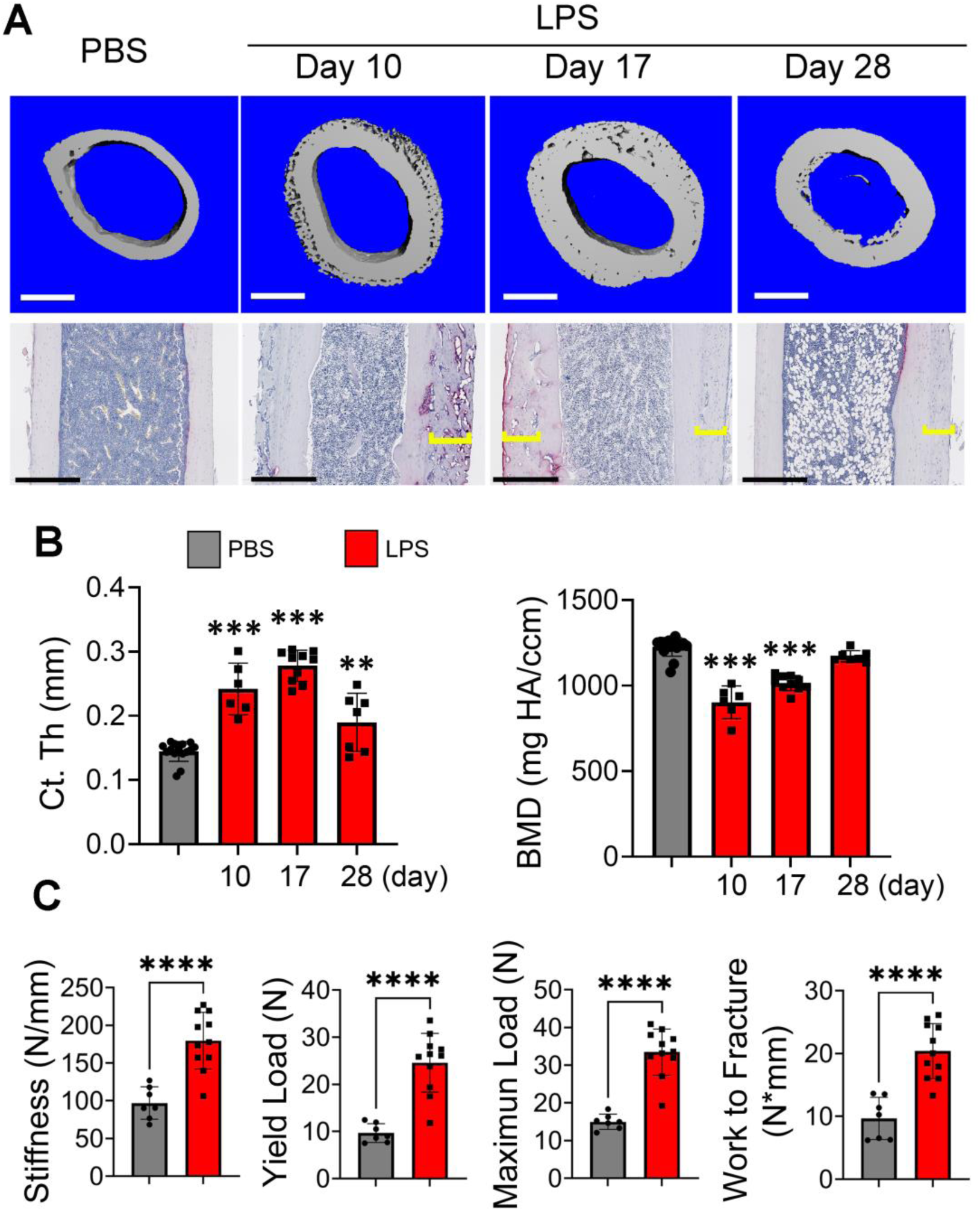
LPS-induced periosteal woven bone undergoes remodeling into mature bone. Three months old WT mice were i.p. injected with PBS (n=15) or 1 mg/kg LPS from *E. coli* (n= 6-10) on day 0, 4, and 8. Mice were sacrificed on day 10, 17, or 28. **A**) Upper panel: Representative μCT images of femoral diaphysis are shown. Scale bar: 500 µm. Lower Panel: Representative femoral sections stained for TRAP activity. Brackets indicate the thickness of the newly formed bone. Scale bar: 400 µm. **B**) Cortical thickness (Ct.Th) and bone mineral density (BMD) analysis at femur midshaft of A. **C**) Three old WT male mice were i.p. injected with PBS or 1 mg/kg LPS from *E. coli* on day 0, 4, and 8. Mice were sacrificed on day 28. Femur biomechanical properties were assessed using a 3-point bending test. **p<0.01; ***p<0.001; ***p<0.0001. One way ANOVA test.

### LPS-induced periosteal bone formation positively correlates with bone marrow cell death

To understand the mechanisms by which LPS induced periosteal bone formation, we analyzed samples from WT mice injected with LPS for different times. The color of bone marrow of LPS- and PBS-treated mice differed, most likely due to inflammation and cell death (Supplementary Fig. 6A, B). Notably, LPS primarily affected bone marrow cells, both CD45⁺ and CD45⁻ populations, whereas periosteal cells and splenocytes were largely unaffected (Supplementary Fig. 6B, C). The reason why CD45⁺ splenocyte populations were unaltered remains unclear. It also remains unknown whether discrete, localized cell death occurs in the periosteum or the adjacent muscle tissues, in addition the bone marrow. Nevertheless, bone marrow of LPS-exposed animals contained numerous leukocytes and displayed a layer of dead cells (arrowheads) interfacing healthy (H) and inflamed (I) areas (Fig. 3A). While morphological changes in bone marrow were visible as early as 5 days after LPS injection, periosteal bone formation was not apparent until 7 days after LPS inoculation (Fig. 3B). µCT analysis confirmed that 7 days following LPS treatment, periosteal bone apposition was confined to the distal metaphysis but had extended to most of the diaphysis by day 10 (Fig. 3C, D). Intriguingly, areas of early periosteal osteogenesis were prominently adjacent to those of compromised bone marrow compared to those of normal bone marrow (Fig. 3E and Supplementary Fig. 4D). These observations were consistent with TUNEL staining, which revealed massive cell death in the interface zone compared to other marrow areas (Fig. 3F and Supplementary Fig. 6E). Collectively, these results suggest that LPS induces bone marrow death, a response that precedes enhancement of cortical thickness.

**Fig. 3.**
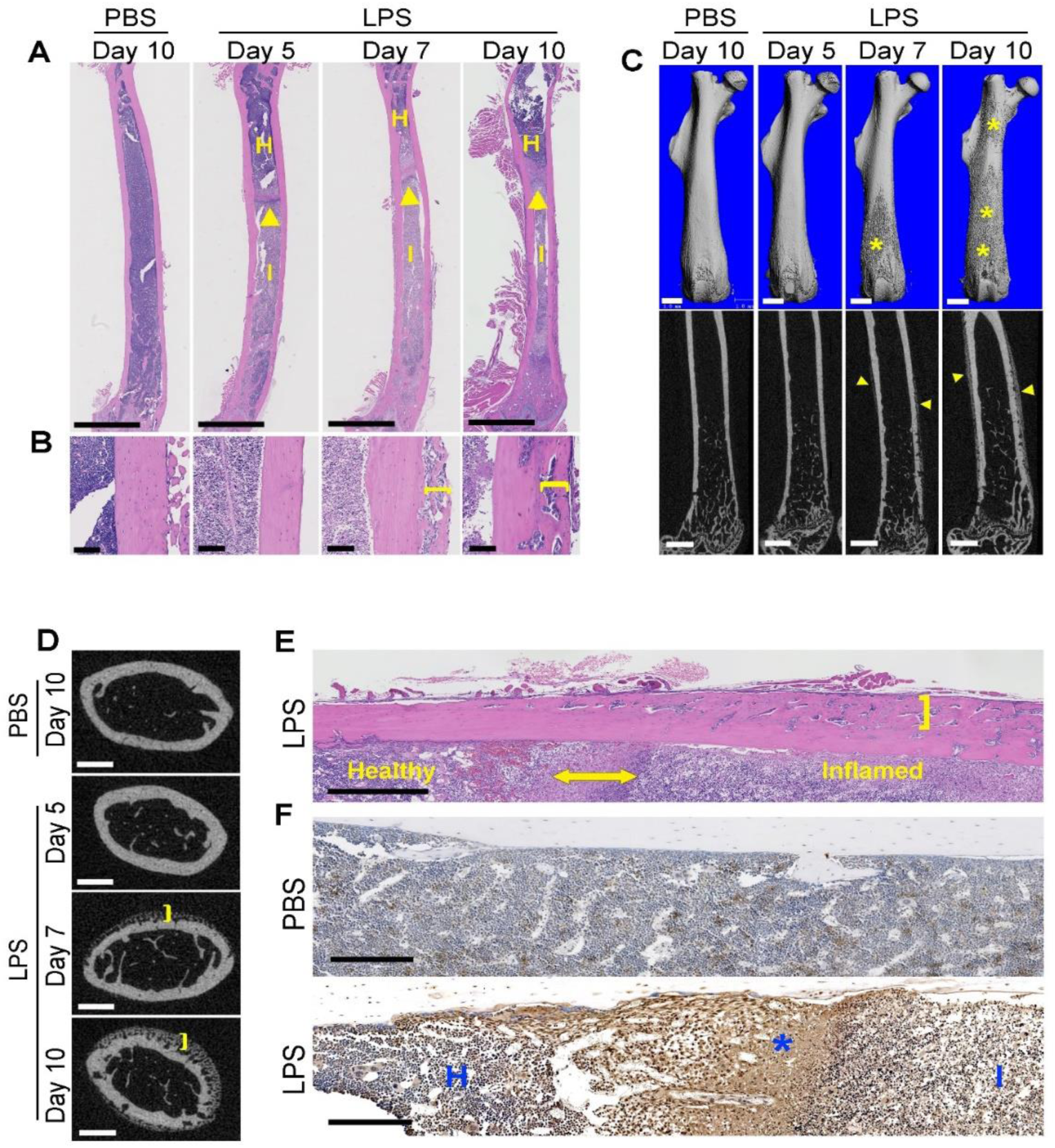
LPS-induced periosteal bone formation positively correlates with bone marrow cell death. Three months old WT mice were i.p. injected with 1 mg/kg LPS or PBS (n = 3 for each group) on day 0, 4, and 8. Mice were sacrificed as indicated. Mice treated with PBS were sacrificed on day 10 as control (PBS). **A**, **B**) Representative images of H&E stained-femoral sections. **A**) Arrowheads show the interface between healthy (H) and inflamed (I) bone marrow. Scale bar: 2 mm. **B**) Brackets indicate the thickness of the newly formed bone. Scale bar: 100 µm. **C**) Representative μCT images of femoral diaphysis. Scale bar: 1mm. Asterisks and arrowhead indicate areas of newlyformed bone. **D**) Representative μCT images of cross section of femora. Scale bar: 500 µm. Brackets indicate areas of newly formed bone. **E**, **F**) Three months old WT mice were i.p. injected with 1 mg/kg LPS (n=7) or PBS (n=5) on day 0, 4 and 8. Mice were sacrificed on day 10. **E**) Representative images of H&E-stained femoral sections. Double ended arrowhead shows the interface between healthy and inflamed pyroptotic bone marrow. Brackets indicate the thickness of the newly formed bone. Scale bar: 500 µm. **F**) TUNEL staining. Staining is stronger in inflamed area (I) compared to heathy area (H) but weaker than in the interface zone (asterisk). Scale bar: 200 µm.

### Non-canonical inflammasome mediates LPS-induced periosteal bone formation

To determine if LPS induced bone marrow pyroptosis and understand the underlying mechanisms, we studied cortical bone outcomes in mice lacking key components of inflammasomes known to be regulated by LPS. These include NLRP3 canonical inflammasome (NLRP3, caspase-1), non-canonical inflammasome (caspase-11), and their shared substrate, GSDMD. Compared to WT mice, loss of NLRP3 or caspase-1 did not affect cortical bone formation (Fig. 4). Lack of IL-1 receptor also failed to prevent LPS effects on bone in all treated mice (Fig. 4 and Supplementary Fig. 7). By contrast, periosteal bone formation was significantly inhibited in mice deficient in *caspase-11*, both *caspase-1* and *caspase-11*, or *Gsdmd* (Fig. 4). Accordingly, LPS-induced bone marrow hypocellularity in WT mice was prevented in *caspase-11^-/-^*and *Gsdmd^-/-^* counterparts (Supplementary Fig. S8). Collectively, these data indicate that LPS primarily activates the non-canonical inflammasome-GSDMD axis to inflict bone marrow pyroptosis, a response that ultimately induces cortical bone formation.

**Fig. 4.**
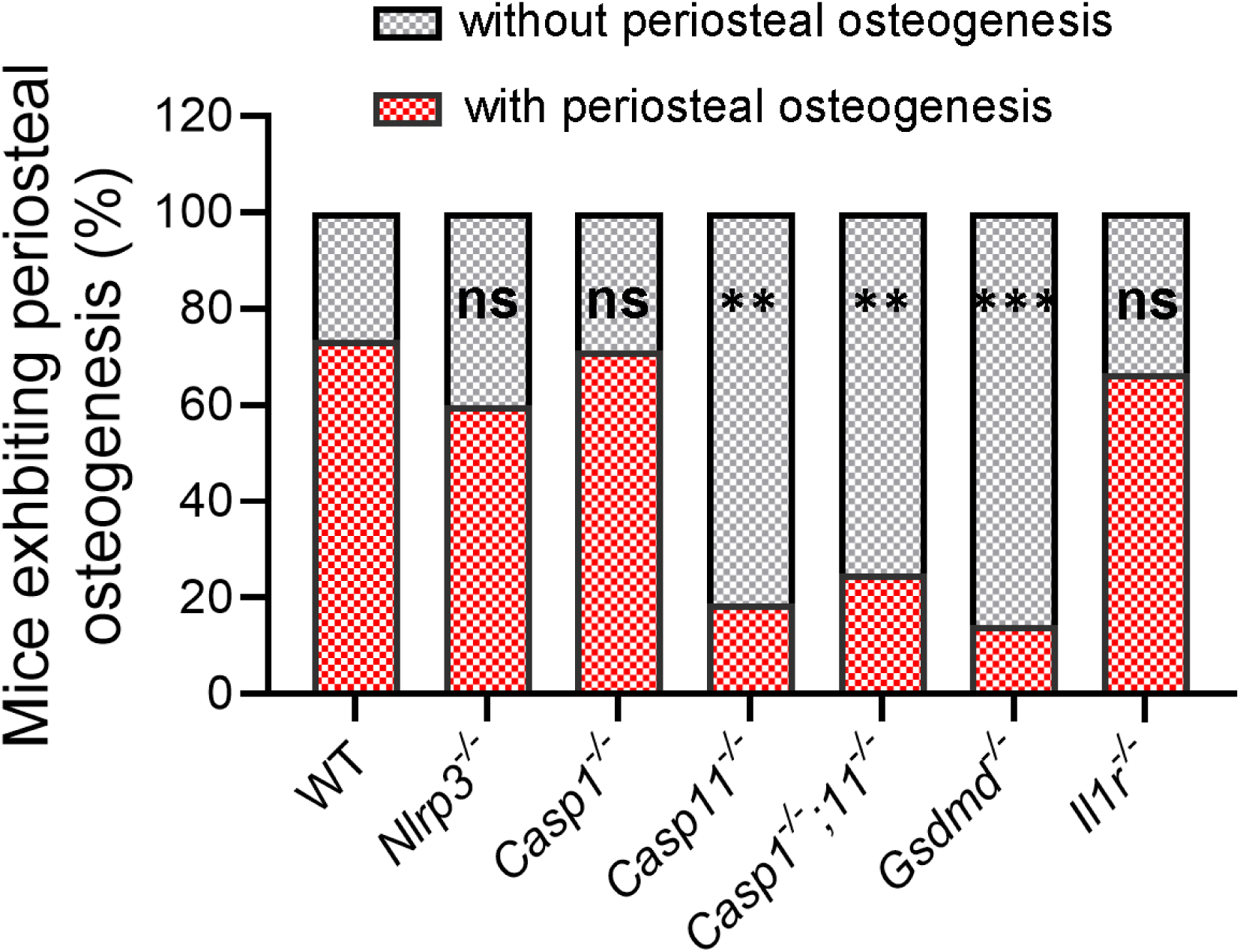
Non-canonical inflammasome mediates LPS-induced periosteal bone formation. Three months old WT and mutant mice were i.p. injected with 1 mg/kg LPS from *E. coli* on day 0, 4 and 8. Mice were sacrificed on day 10 and femurs were collected for µCT analysis. The number of mice exhibiting periosteal bone formation/total number of mice tested is shown under the columns. Chi-Square test vs WT: ns, not significant. **p<0.01, ***p<0.001.

### LPS-induced bone marrow pyroptosis releases osteogenic factors

To reinforce the conclusion that LPS caused bone marrow pyroptosis, we treated mice with PBS or 1 mg/kg LPS for various time points and monitored lactate dehydrogenase (LDH) and IL-1β levels in bone marrow supernatants. LPS induced a time-dependent release of LDH in bone marrow and blood, with significant differences observed at 24 hours and 6 hours post-LPS injection, respectively, compared to PBS controls (Fig. 5A, B). By contrast, LPS caused transient secretion of IL-1β as its levels were sharply increased 6 hours after injection but returned to baseline values by 24 hours.

**Fig. 5.**
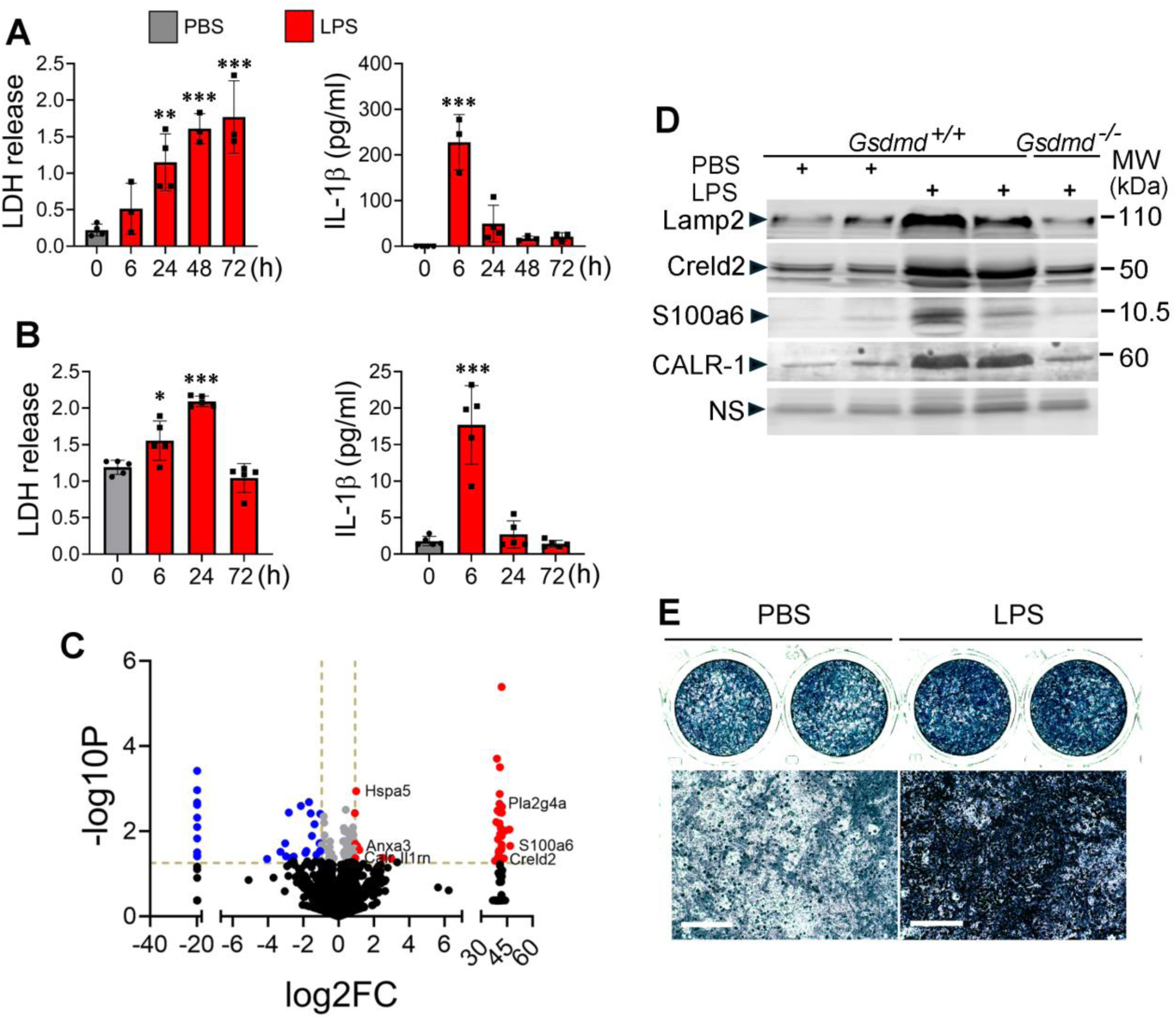
LPS-induced bone marrow pyroptosis releases osteogenic factors. **A, B**) Three-month-old WT male mice were injected with PBS or LPS (n= 3-4) and sacrificed at the indicated time points. Bone marrow supernatants collected from the tibias and femurs centrifuged in tubes containing 100 µl PBS (A) and serum (B) were used for LDH (OD values) and IL-1β analysis. **C**, **D**) Three months old WT mice were i.p. injected with PBS (n=3) or 1 mg/kg LPS (n=3) on day 0 and sacrificed on day 3. Bone marrow was centrifuged, and the supernatants were collected and analyzed by mass spectrometry. **C**) Volcano plot. **D**) Three months old WT and *Gsdmd^-/-^*mice were i.p. injected with PBS or LPS on day 0 and sacrificed on day 3. The supernatants from centrifuged bone marrow were analyzed by immunoblotting. Each lane represents a sample from an individual mouse. **E**) Periosteal cells from WT mice were isolated with collagenase and expanded for 1 week and cultured in osteogenic medium (50 μg/ml of ascorbic acid and 2 mM of glycerol 2-phosphate) for 3 days in the presence of bone marrow supernatants from mice treated with PBS or LPS for 3 days. Cells were stained for alkaline phosphatase activity. Top panels: images of whole wells. Bottom panels: high magnification images. Scale bar: 100 µm. Lamp2, lysosomal-associated membrane protein 2; Creld2, cysteine-rich with EGF-like domains 2; CALR, calreticulin; NS, nonspecific. *p<0.05, ***p<0.001, ****p<0.0001. 2-way ANOVA.

GSDMD pores mediate the secretion of low molecular weight (MW) molecules (MW ≤18 kDa) such as IL-1β and IL-18^49, 50^. Therefore, we hypothesized that the presence in bone marrow fluid of larger cytoplasmic molecules (MW >18 kDa) in response to LPS should be the result of pyroptosis. To test this hypothesis, we employed mass spectrometry to analyze bone marrow supernatants from mice treated with PBS or LPS. Volcano plot showed proteins with increased or decreased levels upon LPS treatment (Fig. 5C). Focusing on the proteins with increased levels, we found that most of them were cytoplasmic resident proteins. They included IL-1 receptor antagonist, Hspa5, calreticulin, S100a6, cPLA2α, Creld2, and annexin 3. To validate mass spectrometry results, we carried out immunoblotting analysis on bone marrow supernatants from mice treated with PBS or LPS, focusing on proteins for which reliable commercial antibodies are available. The levels of Lamp2, Creld2, S100a6, and CALR1 were higher in samples from LPS-exposed mice compared to PBS-treated controls, and this response was blunted in the absence of GSDMD (Fig. 2D). To determine whether fluids containing this cocktail of molecules modulated osteogenesis following their release into the extracellular environment, we collected bone marrow supernatant from WT mice treated with LPS or PBS. Periosteal cells treated with LPS supernatants showed enhanced alkaline phosphatase (ALP) activity and expression of *Alp* and *Col1a1* compared to those treated with PBS supernatants (Fig. 5E and Supplementary Fig. 9). This effect was blunted when the supernatants were pre-treated with proteinase K to digest and inactivate proteins or derived from GSDMD-deficient mice. Comparable results were obtained with osteoprogenitors isolated from bone marrow. Collectively, these findings suggest that the osteogenic activity of LPS supernatants is mediated by proteins released through GSDMD pores.

### Partial depletion of myeloid cells in bone marrow is sufficient to nearly blunt LPS-induced periosteal bone formation

The caspase-11-GSDMD axis is highly active in myeloid cells^51, 52^. To determine the impact of myeloid cell death on LPS-induced periosteal osteogenesis, we mated *Stop-floxed diphtheria toxin A* (*DtA*) mice with *lysozyme M (LysM)-Cre* mice to generate *Stop-Dta*;*LysM-Cre* (*DtA^LysM^*) and control mice. The analysis of the expression LysM (encoded by *lyz2*) in bone marrow by qPCR (Fig. 5A) and FACS (Fig. 5B) confirmed partial ablation of bone marrow myeloid cells. LPS induced periosteal bone formation was significantly reduced in *DtA^LysM^* compared with WT control (Fig. 5C). Thus, LPS-induced death of myeloid cells is involved in periosteal bone outcomes.

### Newly formed periosteal bone induced by LPS is derived from LepR^+^ periosteal SSCs

Bone comprises inner endosteal and outer periosteal compartments with common and likely unique cell populations^11^. Our results showing profound periosteal bone formation in LPS-stimulated mice raise the possibility that SSCs in the periosteum, but not in bone marrow are substantially increased. It has been reported that transgenes driven by 3.6kb Col1a1 promoter (col1*3.6) target both endosteal and periosteal immature OBs^53, 54^. To determine if periosteal and bone marrow SSCs are impacted by LPS, we generated mice in which col1*3.6 drove the expression of herpes simplex virus thymidine kinase (TK) (Tk^Col1*3.6^) to which we administered ganciclovir to deplete SSCs^48, 55^. We found that while depletion of Tk^Col1*3.6^ cells upon ganciclovir administration did not affect LPS-induced bone marrow cell pyroptosis, periosteal bone formation was eliminated in these mice (Fig. 6A), indicating that cortical bone was formed by periosteal SSCs.

**Fig. 6.**
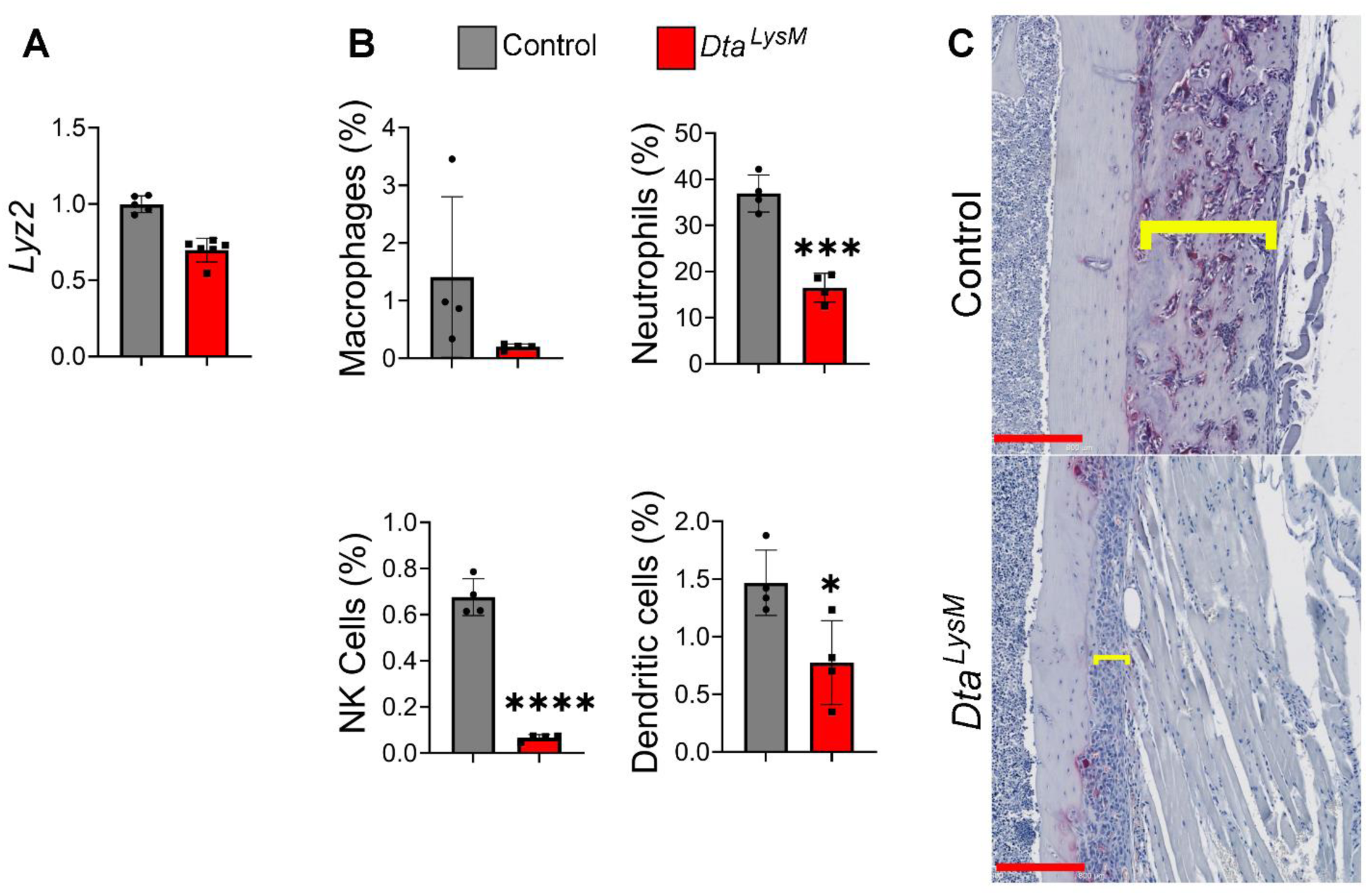
Partial depletion of myeloid cells in bone marrow is sufficient to blunt LPS-induced periosteal bone formation. Three months old WT and *DtA^LysM^*mice were used. Bone marrow cells were used for the analysis of Lyz2 expression by qPCR (**A**) or cells by flow cytometry (**B**). Each black dot represents one mouse. **C**) Representative images of TRAP staining of femoral sections from WT (n= 5) or *DtA^LysM^* (n= 5) mice injected with 1 mg/kg LPS from *E. coli* on day 0, 4, and 8, sacrificed on day 10. Brackets indicate the thickness of the newly formed bone. Scale bar: 200 µm. *p<0.05, ***p<0.001, ****p<0.0001. Unpaired t test.

**Fig. 7.**
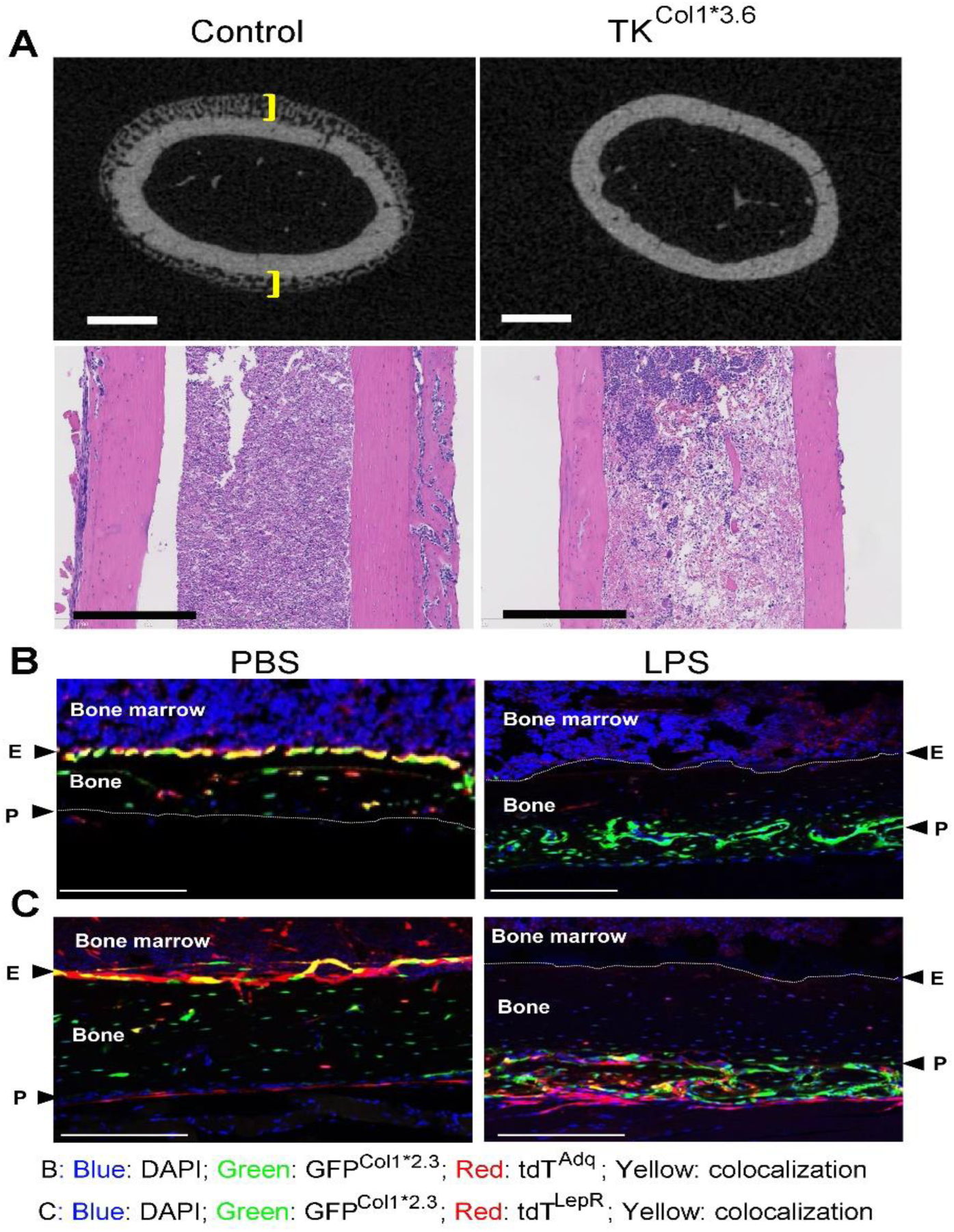
Newly formed periosteal bone induced by LPS is derived from LepR^+^ periosteal SSCs. **A**) Three months old male WT (control; n=11) and Col1*3.6TK mice (n=10) were i.p. injected with 1 mg/kg LPS on day 0, 4, and 8. All mice were i.p. injected with 8 μg/g ganciclovir (GCV), twice daily from day 0 to day 9. Femurs were collected on day 10 and analyzed by µCT and H&E staining. Brackets indicate the thickness of the newly formed bone. Scale: upper panel: 500 µm. Lower panels: H&E staining; scale bar: 400 µm. **B**, **C**) Femurs from 3 months old male *tdT^Adq^*;*GFP^Col1*2.3^* mice (**B**) and *tdT^LepR^*;*GFP^Col1*2.3^* (**C**) mice were i.p. injected with PBS (left panel; n=3) or 1 mg/kg LPS (right panel; n=3) on day 0, 4, and 8 and analyzed on day 10. Representative images are shown. Blue, DAPI; red, tdT^Adq^ (**B**) or tdT^LepR^ (**C**); green, GFP; E, endosteum; P, periosteum. Scale bar: 200 µm.

It has been reported that Adq^+^ SSCs and LepR^+^ SSCs differentiate into OBs both *in vitro* and *in vivo*. To assess the contribution of Adq^+^ and LepR^+^ cells to LPS outcomes, we crossed *stop*- *floxed-tdTomato* (*tdT^fl/+^*), *2.3Col1GFP* (*GFP^Col1*2.3^*), with either *Adq-Cre* (*Adq^Cre^*) or *Lepr-Cre* (*Lepr^Cre^*) mice to respectively generate *Adq^Cre^;tdT;GFP^Col1*2.3^* (*tdT^Adq^*) mice and *Lepr^Cre^;tdT;GFP^Col1*2.3^* (*tdT^LepR^*) mice and monitored the colocalization of labelled cells. 2.3Col1 was used as a marker of mature OBs^53, 56^. In PBS treated-mice, while *tdT^Adq^* only labels endosteum (Fig. 6B, left panel), *tdT^lepR^* targets both endosteum and periosteum (Fig. 6C, left panel). After LPS stimulation, *tdT^Adq^*did not target any GFP^Col1*2.3+^ OBs in the newly formed periosteal bone (Fig. 6B, right panel), but *tdT^LepR^* targeted approximately 15% GFP ^Col1*2.3+^ OBs in this new tissue (Fig. 6C, right panel; Supplementary Fig. 10). This suggests that the newly formed periosteal OBs are not derived from Adq^+^ cells, but LepR^+^ cells contribute, at least in part, to periosteal bone formation following LPS stimulation.

## Discussion

LPS primes canonical inflammasomes, such as those formed by NLRP3, and directly triggers noncanonical inflammasomes, resulting in the release of IL-1β and IL-18 or induction of pyroptosis ^33–36^. In this study, we unexpectedly found that low systemic levels of LPS not only caused inflammation and bone resorption as expected but also led to significant periosteal bone accrual, which underwent remodeling and integrates with the existing bone, resulting in a stronger overall bone structure compared to its pre-LPS state. Using various genetic mouse models, we showed that this periosteal response was associated with pyroptosis of both CD45^+^ cells and CD45^-^ cells in the bone marrow driven by the caspase-11-GSDMD pathway. The exact reason why the LPS regimen used in this study induced pyroptosis and cortical bone response remains unclear. However, a recent meta-analysis highlighted that the effects of LPS on trabecular and cortical bone structures varied depending on the dose, route of administration, and duration of exposure^42^. Given the complexity of LPS’s effects, further research is needed to better understand the mechanisms driving the responses observed in this study.

The release of intracellular contents into the extracellular space is a critical step in the inflammatory responses induced by lytic cell death. In addition to IL-1β, IL-18, and various other proteins, pyroptotic cells also released metabolites and eicosanoids like PGE2, which were shown to promote wound healing^57, 58^. Moreover, remnants of pyroptotic cells, often crowned with F-actin-rich filopodia, have been suggested as a mechanism for initiating adaptive immune responses^58^. Our findings revealed that LPS-induced pyroptosis in bone marrow cells led to the release of several proteins, most of which were resident intracellular macromolecules. We confirmed that bone marrow supernatants from *Gsdmd^+/+^*mice, but not *Gsdmd^-/-^* littermates treated with LPS, enhanced OB differentiation *in vitro* when compared to supernatants from PBS-treated mice. Collectively, these findings highlight a novel mechanism in which pyroptotic myeloid cells release molecules that directly or indirectly promote the differentiation of neighboring mesenchymal cells. Thus, in addition to its established roles in limiting replicative niches during infection, propagating inflammation, promoting tissue repair, and triggering adaptive immune responses^57, 58^, pyroptosis also plays a pivotal role in facilitating cell differentiation.

Osteogenesis is initiated by the proliferation and differentiation of SSCs. While the mesenchymal nature of these cells is well established, their precise identification remains challenging. Bone marrow Adq^+^ cells have the potential to differentiate into OBs, and within adult bone marrow, LepR^+^ SSCs are the primary source of new OBs for bone maintenance. Although Adq^+^ and LepR^+^ cells almost entirely overlap in bone marrow, periosteal cells are predominantly targeted by LepR^+^ cells, rather than Adq^+^ cells. Our findings suggest that bone formation occurred in the periosteum but not in the endosteum, likely because osteoprogenitors survived in the periosteal envelope but were rapidly lost in the bone cavity marrow following LPS exposure. As pyroptosis-driven inflammation gradually resolves, trabecular bone formation is subsequently initiated. Lineage tracing data further confirmed that tdT^LepR^ cells, but not tdT^Adq^ cells, targeted OBs in the newly formed periosteum. These results suggest that: 1) newly differentiated periosteal OBs did not originate from Adq^+^ cells, and LepR^+^ cells were at least partially responsible for the periosteal bone formation induced by LPS; 2) bone marrow Adq^+^ cells were unable to migrate into the periosteum, a view that is consistent with the massive cell death in the bone marrow compartment.

LPS induced the death of CD45^-^ cells in the bone marrow, but not in the periosteum. Similarly, it led to the loss of tdT⁺LepR⁺ cells in the bone marrow while sparing those in the periosteum. In addition, SSCs from both compartments exhibited enhanced *in vitro* osteogenic potential when exposed to supernatants from LPS-treated mice. Together, these findings support our working model that LPS promotes bone formation in the periosteum, but not in the endosteum, likely due to compartment-specific differences in SSC survival. However, several critical questions remain to be addressed to fully support this model. First, do macromolecules released by pyroptotic cells influence bystander cells extracellularly, or is internalization required? Precedent for extracellular activity of intracellular macromolecules exists; for example, histones and DNA can propagate inflammation when released, as seen with macrophage and neutrophil extracellular traps⁽⁶⁵⁻⁶⁷⁾. Second, how do factors released from pyroptotic bone marrow cells traverse the cortical bone to reach periosteal osteoprogenitors? In this context, might these macromolecules be transported via transcortical blood vessels²⁹,³⁰ and/or act on osteocytes near the endosteum, initiating signaling through the lacunar-canalicular networks³¹,³² to promote periosteal bone formation? Despite these unresolved questions, our study underscores the concept that pyroptosis is not merely an endpoint, but rather a dynamic response that triggers downstream cascades. Indeed, we report that the occurrence of pyroptosis in bone marrow myeloid cells promotes not only the expected inflammation-associated osteoclastogenesis but also osteogenesis.

## Materials and Methods

### Mice and Reagents

*iDtr* (*Lox-Stop-Lox-ROSA-DTR*) mice, *Rosa-DtA* (*Lox-Stop-Lox-ROSA-DTA*) mice, *Col1a1*2.3GFP* mice, *Adipoq-Cre* mice, *LepR-Cre* mice, *LysM-Cre* mice, *tdTomato* (Ai9) mice, *Nlrp3^-/-^* mice, *Casp1^-/-^* mice*, Casp11^-/-^* mice*, Casp1^-/-^;11^-/-^* mice, and *Gsdmd^-/-^* mice were purchased from The Jackson Laboratory. *TK-3.6Col1a1* mice [^55^] were kindly provided by Robert Jilka and Charles O’Brien (University of Arkansas). All animals were housed in the specific pathogen free (SPF) animal facility of Washington University School of Medicine, where they were maintained at 22°C in a 12-hour light-dark cycle. Animal work was performed according to the policies of IACUC at Washington University School of Medicine in St. Louis. Mice were analyzed under approved IACUC protocols.

Mice were crossed in-house to generate *Adipoq^Cre^;Dtr* (*Dtr^Adq^*) mice, *Lepr^Cre^;Dtr* (*Dtr^Lepr^*), *Adipoq^Cre^;Col1*2.3GFP;Ai9* (*tdT^Adq^*;*GFP^Col*2.3^*) and *LepR^Cre^*;*Col1*2.3GFP;Ai9* (*tdT^Lepr^*;*GFP^Col*2.3^*) on a C57Bl/6 background. To ablate Cre-positive cells, diphtheria toxin (Biological List laboratories, Cat #150) (100 ng/mouse/day) was daily i.p. injected into mice. Ganciclovir (Acros Organics, Cat# 461710050) was injected at 8 μg/g i.p. twice daily in saline. LPS (Sigma Aldrich, L2630) was given at 1 mg/Kg i.p. as indicated. LPS, collagenase I, and collagenase IV were purchased from Sigma.

### Microcomputed tomography (μCT)

The whole femur was scanned using μCT50 scanner (Scanco Medical AG, Bassersdorf, Switzerland; 70 kVp, 57 µA, 700 ms integration time, 7.4 µm voxel size). A lower threshold of 220 was used for evaluation of all scans.

### Biomechanical testing

Three old WT male mice were i.p. injected with PBS or 1 mg/kg LPS from *E. coli* on day 0, 4, and 8. Mice were sacrificed on day 28. Femur biomechanical properties were assessed using a 3- point bending test as previously described^37, 59^. Briefly, femurs were dissected and cleaned of all soft tissue. The ends of each femur were potted using polymethylmethacrylate (PMMA) in 6mm- diameter ×12-mm-length acrylic tubes. The bone was centered using a custom fixture, leaving approximately 4.2 mm of exposed bone between potting tubes. All samples were wrapped in PBS-soaked gauze to preserve hydration while the PMMA cured overnight. The following day, each sample was loaded into a custom-built torsion machine with a 25 inch-ounce load cell controlled with LabVIEW software (LabVIEW 2014; National Instruments, Austin, TX, USA). The machine held one of the potted femoral ends in a fixed position while rotating the other potted tube at 1 degree/s until fracture. The stiffness (N/mm), Yield force (N), work to fracture (N*mm) and maximum load (N) were calculated from the resulting torque-rotation graphs (Matlab; MathWorks, Natick, MA, USA).

### Histology and histomorphometry

Femurs and tibias were fixed in 10% neutral buffered formalin, followed by decalcification in 14% EDTA for 10 days, paraffin embedding, and TRAP or H&E staining.

### Microscopic analysis

Bones were fixed with 10% neutral buffered formalin 24-48 hours at room temperature and decalcified in 14% EDTA for 3 days. The bones were then infiltrated with 30% sucrose overnight at 4 °C. 10-µm thick cryoprotection sections were mounted with antifade mounting medium with DAPI (Vector Laboratories), and images were acquired with a confocal microscope (Nikon C-1 confocal system).

### TUNEL staining

ApopTag Peroxidase in Situ Apoptosis Detection Kit was purchased from Sigma. Paraffin sections (4 µm thick) were subjected to TUNEL staining according to the manufacturer’s instructions and analyzed using ImageJ. The percentage of TUNEL⁺ cells was determined by dividing the number of TUNEL⁺ cells by the total number of cells.

### Reporter bone analysis

Femurs of *tdT^lepR^;GFP^Col1*2.3^* or *tdT^Adq^; GFP^Col1*2.3^* were fixed in 10% neutral buffered formalin, followed by decalcification in 14% EDTA for 3 days. The bones were then infiltrated with 30% sucrose overnight at 4°C for cryoprotection and embedded in optimal cutting temperature (Tissue-Tek). Sections of 10-μm thickness were prepared with a Leica cryostat equipped with Cryojane (Leica, IL). tdT^+^ and GFP^+^ cells were analyzed using ImageJ.

### Bone marrow supernatant and blood collection and ELISA analysis

After removal both sides of long bones, bone marrow was centrifuged at 3000 rpm into PBS (100µl/mouse) and supernatants were kept at -80 °C. Blood was drawn from submandibular vein with BD Microtainer collection tubes. Cell death was assessed by detection of LDH in marrow supernatant and serum using the LDH Cytotoxicity Detection Kit (Takara, CA). IL-1β levels in marrow supernatant and serum were measured by an ELISA kit (eBioscience, NY).

### Proteomics analysis

Mouse bone marrow was centrifuged in a tube containing 100 µl of PBS. Collected samples were used for proteomics analysis by the Mass Spectrometry Technology Access Center at McDonnell Genome Institute (MTAC@MGI) at Washington University School of Medicine. Briefly, after measuring protein concentrations, each sample was reduced, alkylated, and digested with trypsin according to the optimized MTAC’s SOP. Digested peptides were desalted on C18 spin columns, quantified and the same normalized amount of peptides were analyzed by LC-MS/MS using an Orbitrap instrument online with nanoUHPLC. Data were searched against a mouse database using MaxQuant search engine and subjected to label-free quantification (LFQ) based on the MS1 peptide intensity. Proteins were filtered for >1 unique peptide. For statistical analysis, the LFQ intensities will be Log2 transformed and grouped into biological groups. Imputation was applied to fill in missing values based on normal distribution, which is recommended for LFQ. Pair-wise t test with FDR correction was employed to determine significant differences among the groups.

### Western blot analysis

Three months old WT and *Gsdmd^-/-^* mice were i.p. injected with PBS or LPS on day 0 and sacrificed on day 3. The supernatants from centrifuged bone marrow were analyzed by immunoblotting. Protein concentrations were determined by the Bio-Rad Laboratories method, and equal amounts of proteins were subjected to SDS–polyacrylamide gel electrophoresis using 12% or 15% gels. Proteins were transferred onto nitrocellulose membranes and incubated with antibodies against Lamp2 (DSHB, UIowa, IA), Creld2 (DSHB, UIowa, IA), S100a6 (DSHB, UIowa, IA), CALR-1(DSHB, UIowa, IA) overnight at 4°C followed by incubation at room temperature for 1 hour with secondary goat anti-mouse IRDye 800 (Thermo Fisher Scientific, MA), goat anti-rat Alexa Fluor 680 (Molecular Probes, Oregon) or goat anti-rabbit Alexa Fluor 680 (Thermo Fisher Scientific, MA), respectively. The results were visualized using the Odyssey Infrared Imaging System (LI-COR Biosciences, NE).

### Isolation of bone marrow cells and periosteal cells and osteoblast differentiation

Bone marrow cells and periosteal cells were isolated with collagenases digestion as previously described^60, 61^. After 1 week expansion in αMEM with 20% FBS, 5x10^4^ cells/well/ml were seeded into 24 well plate for 3 days after which osteoblast differentiation medium containing 50 μg/ml ascorbic acid and 2 mM β-glycerophosphate in αMEM was added. Cells were stained for alkaline phosphatase activity.

### Flow cytometry analysis

Cells were washed with sterile DPBS and pelleted by centrifugation at 500 × g for 5 minutes at 4 °C. The cell pellets were resuspended in 50 μl Zombie NIR viability dye (BioLegend, 423105; 1:1000 in PBS) and incubated on ice for 5 minutes. After washing with FACS buffer, cells were pelleted and incubated with 25 μl Fc block (BD Biosciences, 553141; 1:100 in Brilliant Stain Buffer) for 15 minutes on ice. Cells were then stained by adding 25 μl of a 2× cocktail of fluorophore-conjugated primary antibodies directly to the suspension. The 2X primary antibody cocktail is comprised of rat anti-mouse SiglecF-BV421 (clone E50-2440, BD Horizon, 562681), rat anti-mouse/human CD11b-Pacific Blue (clone M1/70, BioLegend, 101224), rat anti-mouse F4/80-BV650 (clone BM8, BioLegend, 123149), rat anti-mouse Ly6G-BV711 (clone 1A8, BioLegend, 127643), rat anti-mouse CD45-PerCP (clone 30-F11, BioLegend, 103130), mouse anti-mouse CD64-PE/Dazzle594 (clone X54-5/7.1, BioLegend, 139320), Armenian hamster anti-mouse CD11c-PE/Cy5.5 (clone N418, Invitrogen/eBioscience, 35-0114-82), rat anti-mouse MHC- II-PE/Fire 810 (clone M5/114.15.2, BioLegend, 107667). Following a 30-minute incubation on ice, cells were washed twice with FACS buffer and resuspended in 200 μl FACS buffer. An aliquot of 100 μl was immediately analyzed live on a Cytek Aurora spectral flow cytometer.

### Data Representation and Statistical Analysis

Data are expressed as mean ± SD. Statistical significance was determined using unpaired Student’s 2 tailed t test, 2-way ANOVA test or chi-square test. *p<0.05, **p< 0.01, ***p<0.001 in all experiments.

## Acknowledgements

This research was supported by the following NIH grants to GM (R01 AR076758, R01 AG077732, and R01 AI161022) and STL (P30 AR074992, R37 AR046523, R01 DK111389, R01 CA258325), grant from Department of Defense PR200609 to SLT, The Foundation for Barnes-Jewish Hospital to SLT, and TRPA funding from Department of Pathology and Immunology, AMP division to SLT and WZ, and American Heart Association Postdoctoral Fellowship 24POST1244220 to WJ. Mass Spectrometry analyses were performed by the Mass Spectrometry Technology Access Center at McDonnell Genome Institute (MTAC@MGI) at Washington University School of Medicine. We thank Washington University Musculoskeletal Research Center (supported by P30-AR074992 and S10 OD028573) for the assistance with histology service and µCT scan.

