## Supplemental figures for "Activation of the noncanonical inflammasome-GSDMD pathway triggers pyroptosis in bone marrow and promotes periosteal bone formation"

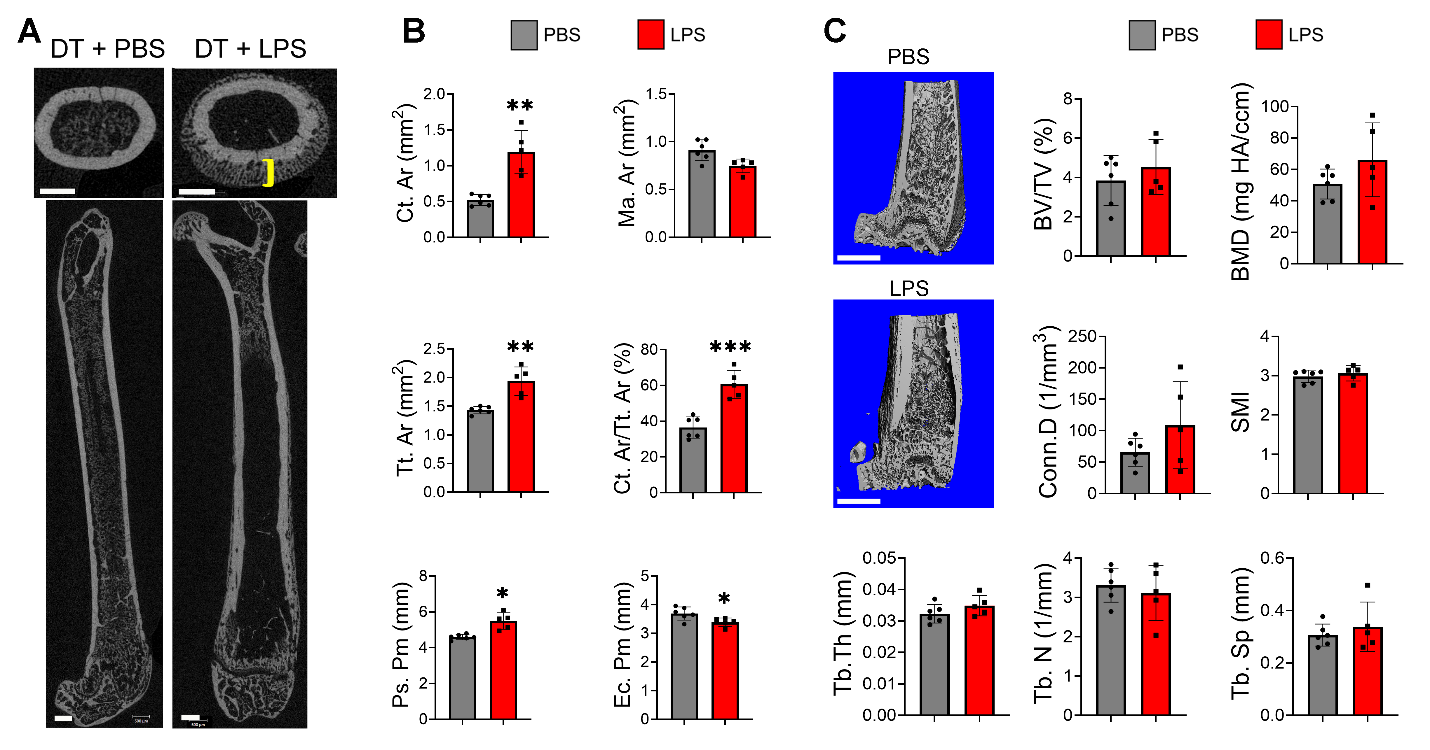

**Supplementary Fig. 1. LPS stimulates periosteal bone formation while endosteal bone formation is unaffected**. **A**) Three months old male *Dtr^Adq^* mice were i.p. injected with DT (100 ng/mouse) once daily from day 0 to day 9 and 1 mg/kg LPS (n=10) or PBS (n=13) on day 0, 4, and 8. Femurs were collected on day 10 and analyzed by µCT. Scale bar: 500 µm. **B, C**) Three months old WT mice were i.p. injected with PBS (n=6) or 1 mg/kg LPS (n=5) on day 0, 4 and 8, mice were sacrificed on day 10. μCT analysis of cortical bone (**B**) or trabecular none (**C**). Representative images (**A, C**). Brackets indicate the thickness of the newly formed bone. *p<0.05, **p<0.01. Unpaired t-test.

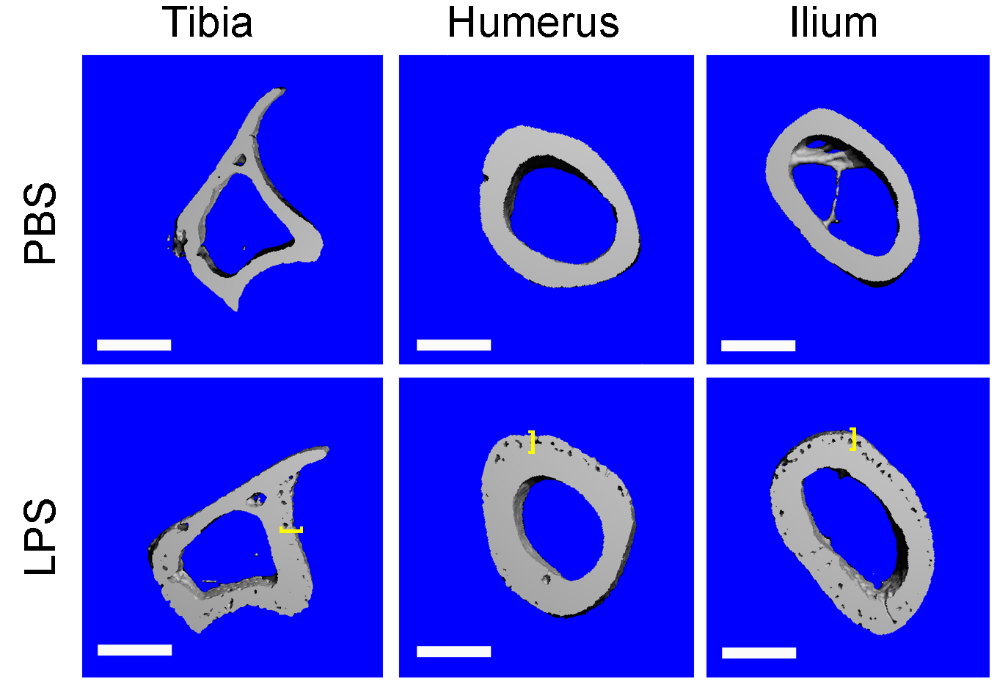

**Supplementary Fig. 2.** **LPS stimulates periosteal bone formation at multiples**. **A**-**F**) Three months old WT mice were i.p. injected with PBS (n=5) or 1 mg/kg LPS from *E. coli* (n=6) on day 0, 4, and 8. Mice were sacrificed on day 10. Representative μCT images are shown. Scale bar: 500 µm.

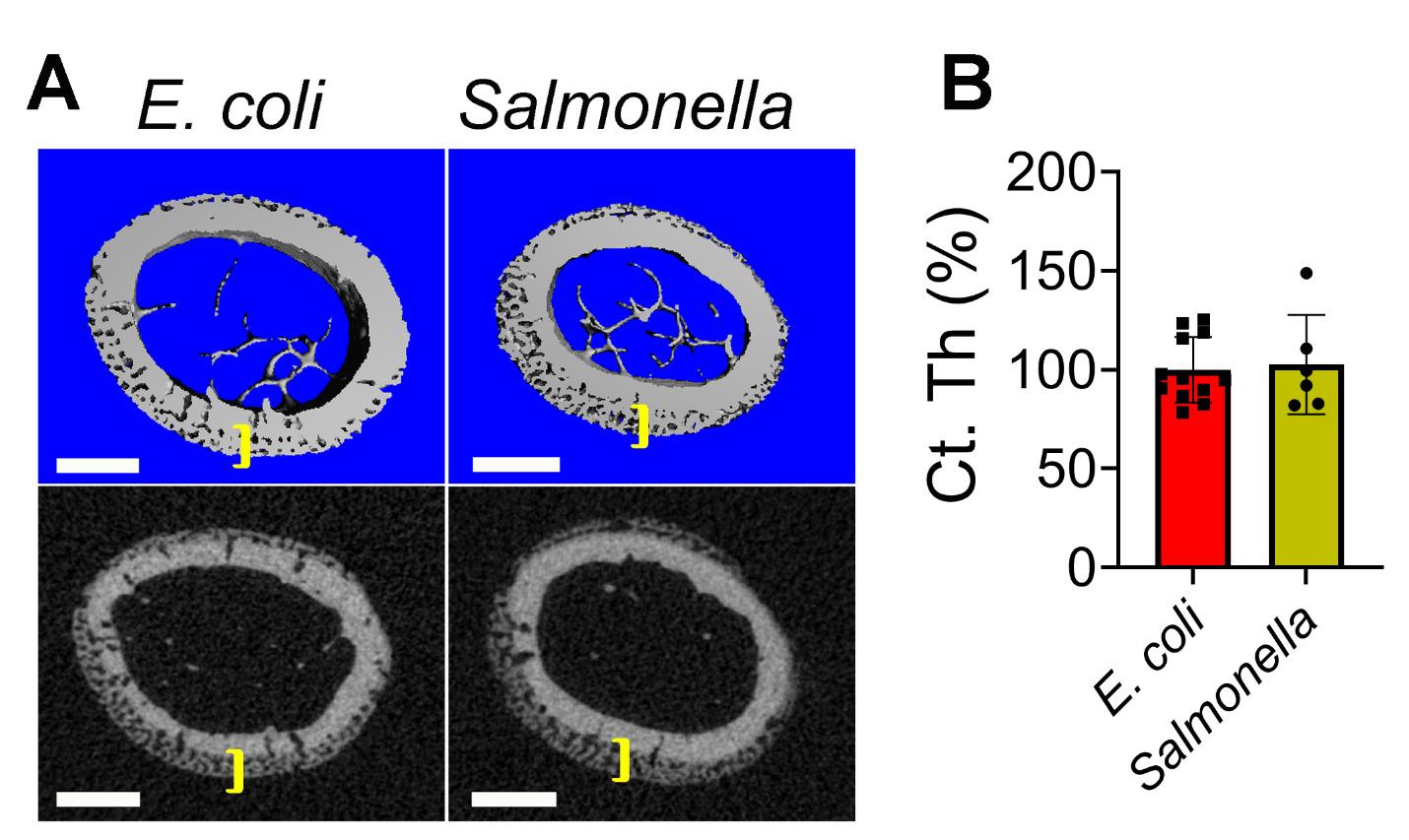

**Supplementary Fig. 3.** **LPS from *E. coli* and *Salmonella* similarly stimulates periosteal bone formation**. Three months Three months old WT mice were i.p. injected with 1 mg/kg LPS from *E. coli* (n=12) or 1 mg/kg LPS from *Salmonella* (n=6) on day 0, 4, and 8. Mice were sacrificed on day 10, and femurs were analyzed by µCT. **A**) Representative images. Brackets indicate the thickness of the newly formed bone. Scale bar: 500 µm. **B**) Percentage of cortical thickness (Ct.Th) changes. *E. coli*’s LPS was set as 100% in **D**). *p<0.05, **p<0.01. Unpaired t-test.

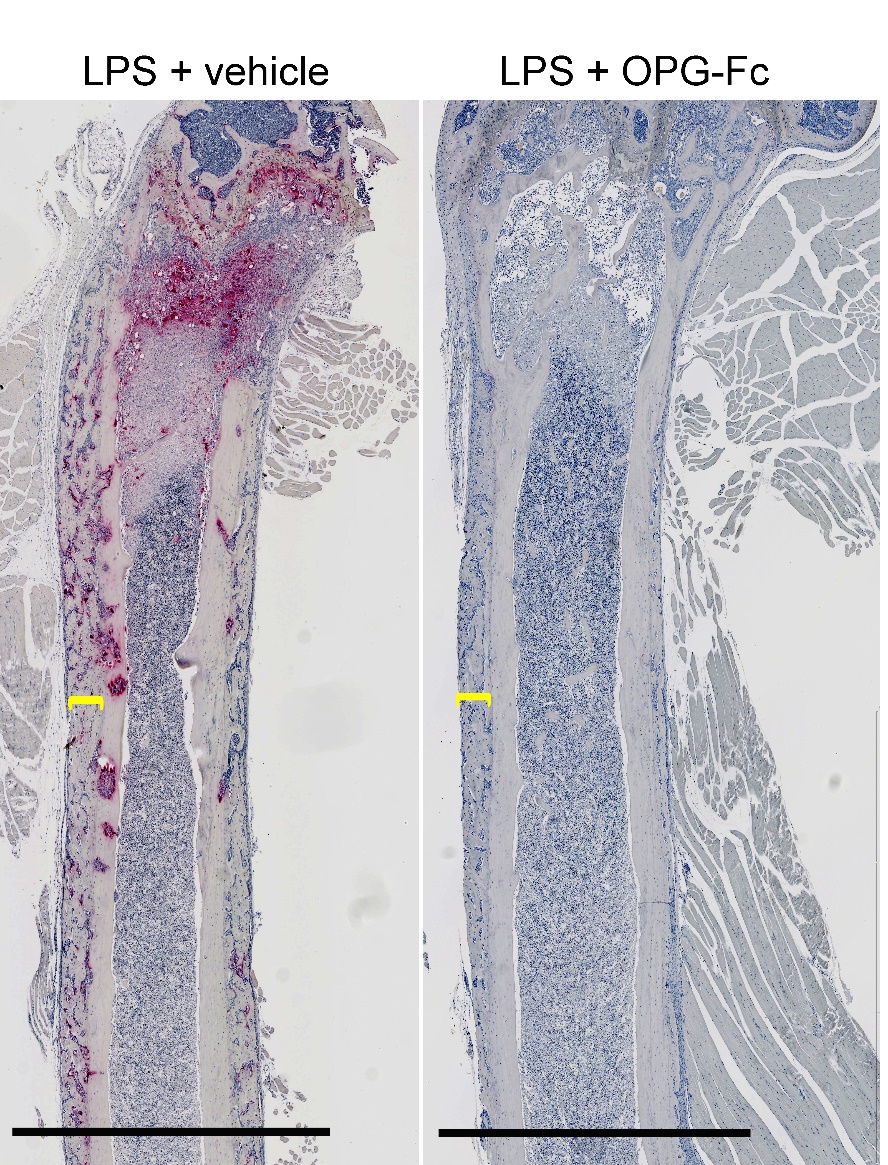

**Supplementary Fig. 4. LPS-induced periosteal bone formation is decoupled from bone resorption.** Three months old WT mice were i.p. injected with 1 mg/kg LPS with (n=10) or without OPG-Fc (5 mg/kg/mouse; n=10) on day 0, 4, and 8. Mice were sacrificed on day 10 and femurs were used for TRAP staining. Representative images are shown. Brackets indicate the thickness of the newly formed bone. Scale bar: 500 µm. Scale bar: 2 mm.

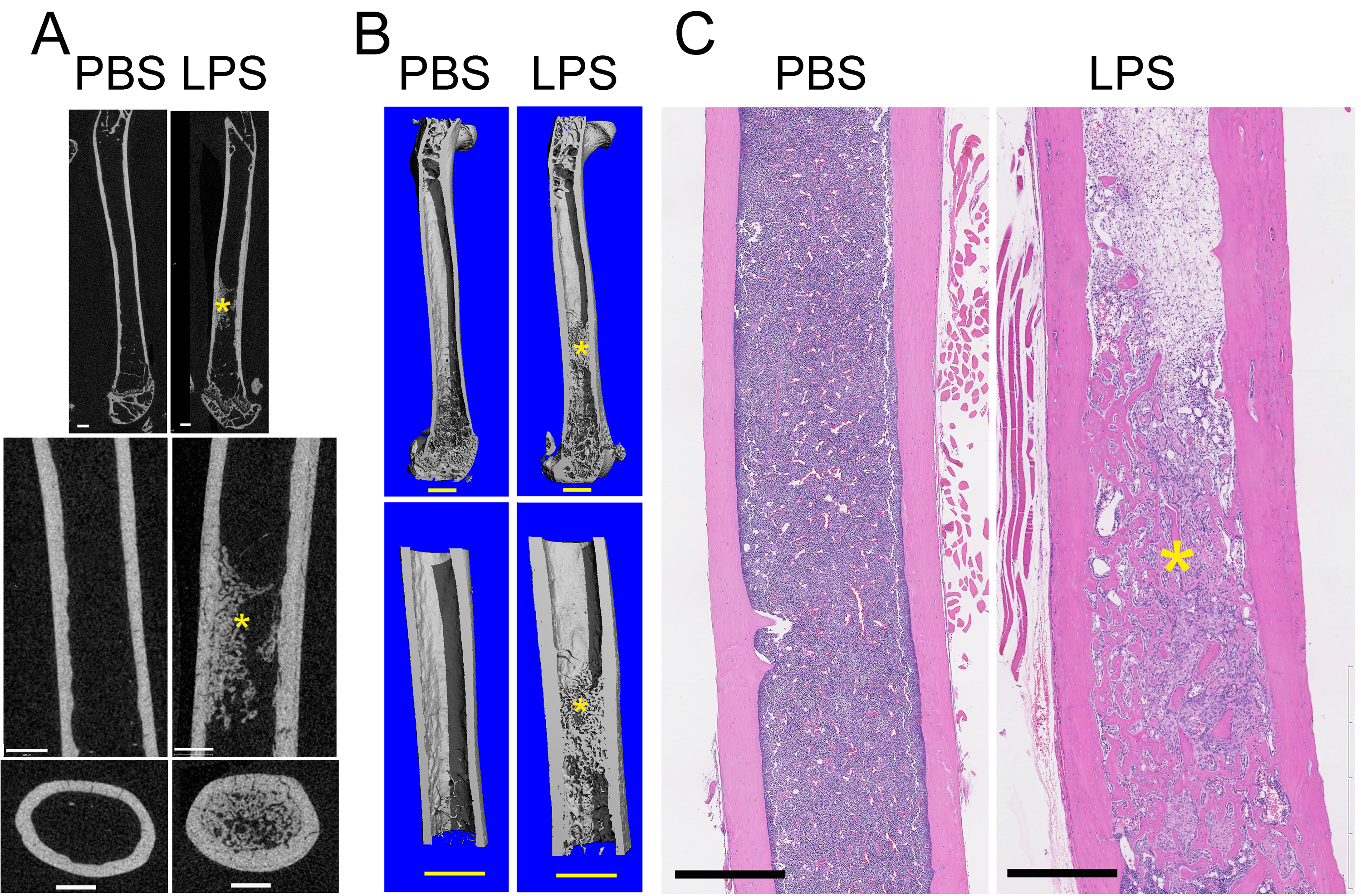

**Supplementary Fig. 5.** **LPS-treated mice subsequently exhibit increased bone trabeculation.** Three months Three months old WT mice were i.p. injected with PBS or 1 mg/kg LPS from *E. coli* (n=7) on day 0, 4, and 8. Mice were sacrificed on day 28, and femurs were analyzed by µCT or histology. Representative images are shown. Asterisks indicate the areas of bone trabeculation. Scale bar: 500 µm.

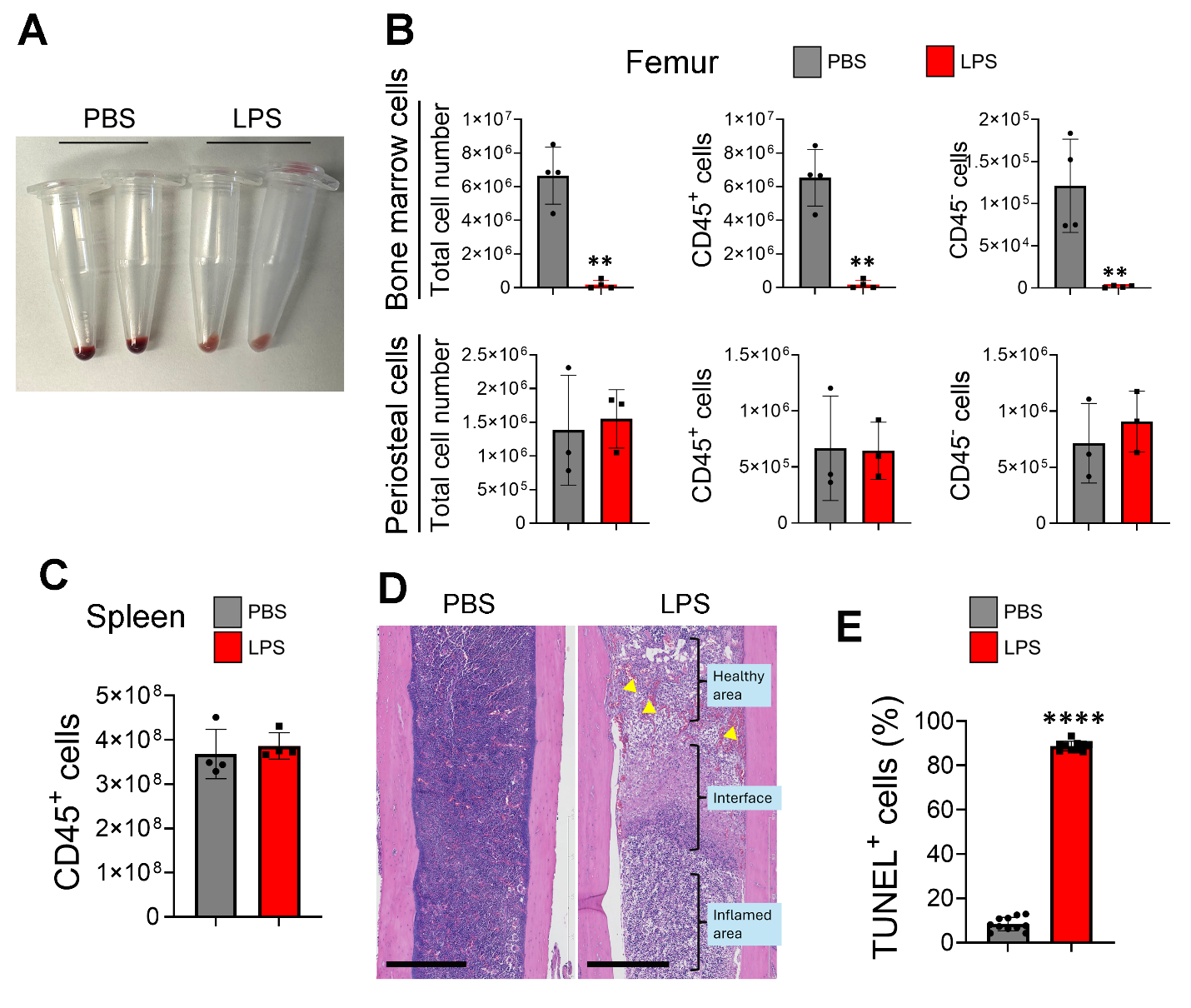

**Supplementary Fig. 6. LPS causes inflammation and the death of bone marrow cells, but not periosteal cells and splenocytes**. **A**) Three months old WT male mice were i.p. injected with PBS (n=3) or 1 mg/kg LPS (n=3) on day 0 and sacrificed on day 3. **A**) Representative pictures of centrifuged bone marrow. Flow cytometry analysis of bone marrow cells, periosteal cells (**B**), and splenocytes (**C**). **D**) Three months old WT male mice were i.p. injected with PBS (n=5) or 1 mg/kg LPS (n=7) on day 0, 4, and 8. Mice were sacrificed on day 10 and the femurs were used for H&E staining. Representative images showing numerous blood vessels in healthy bone marrow area of LPS-injected mice (arrowheads); the interface zone includes dead cells (absence of nuclei), and inflamed marrow area is densely populated by neutrophils. Scale bar: 400 μm. E) Quantitative data of TUNEL^+^ cells. ****p<0.0001.

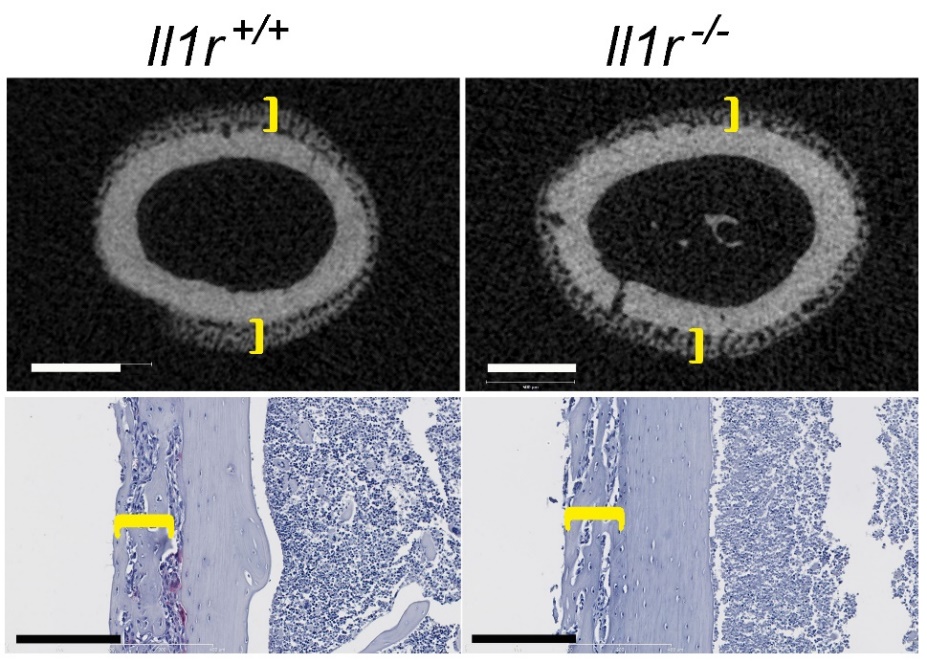

**Supplementary Fig. 7. Loss of IL-1 signaling does not prevent LPS-induced periosteal bone formation**. Three months old *Il1r^+/+^* (n= 6) and *Il1r^-/-^* mice (n=6) were i.p. injected with 1 mg/kg LPS on day 0, 4 and 8. Mice were sacrificed on day 10 and femurs were used for µCT and TRAP staining. Representative images are shown. Brackets indicate the thickness of the newly formed bone. Scale bar: 500 µm (upper panels) and 200 µm (lower panels).

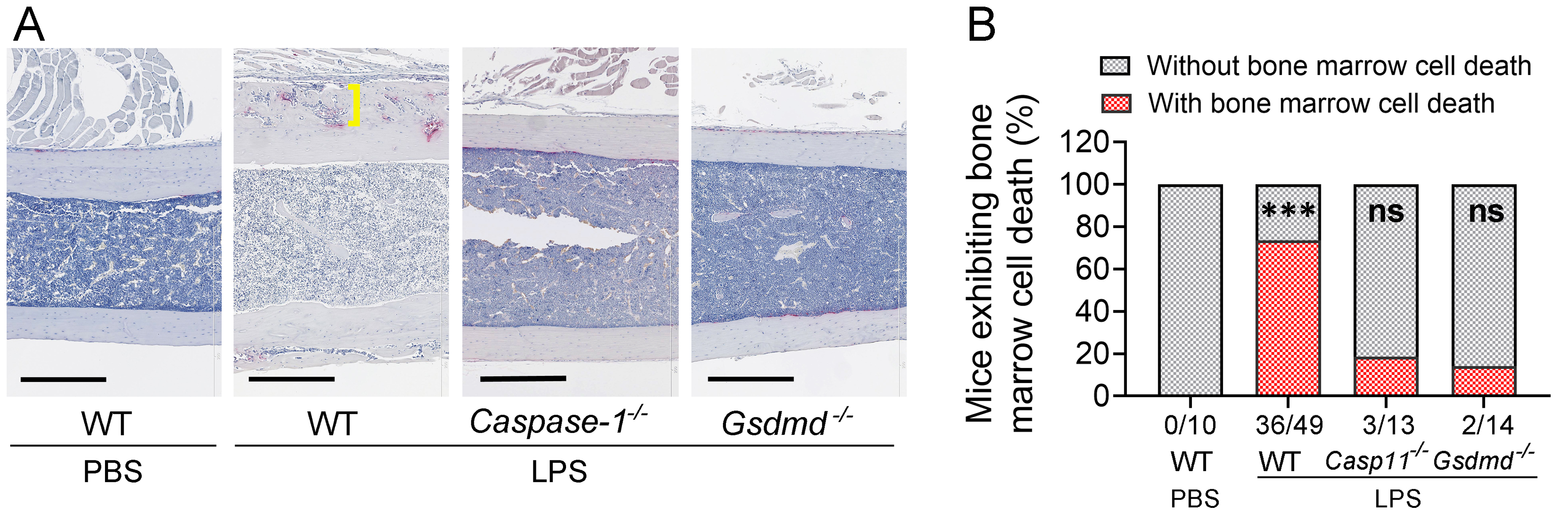

**Supplementary Fig. 8. LPS-induced periosteal bone formation and bone marrow hypocellularity is prevented in mice lacking caspase-11 or GSDMD**. Three months old WT mice (n=10), *caspase-11^-/-^* mice (n=13) and *Gsdmd^-/-^* mice (n=14) mice were i.p. injected with PBS or 1 mg/kg LPS from *E. coli* on day 0, 4 and 8. Mice were sacrificed on day 10. **A**) Femoral sections were stained for TRAP activity. **B**) Sections were visualized under the microscope for evidence of hypocellularity. Brackets indicate the thickness of the newly formed bone. Scale bar: 400 µm.

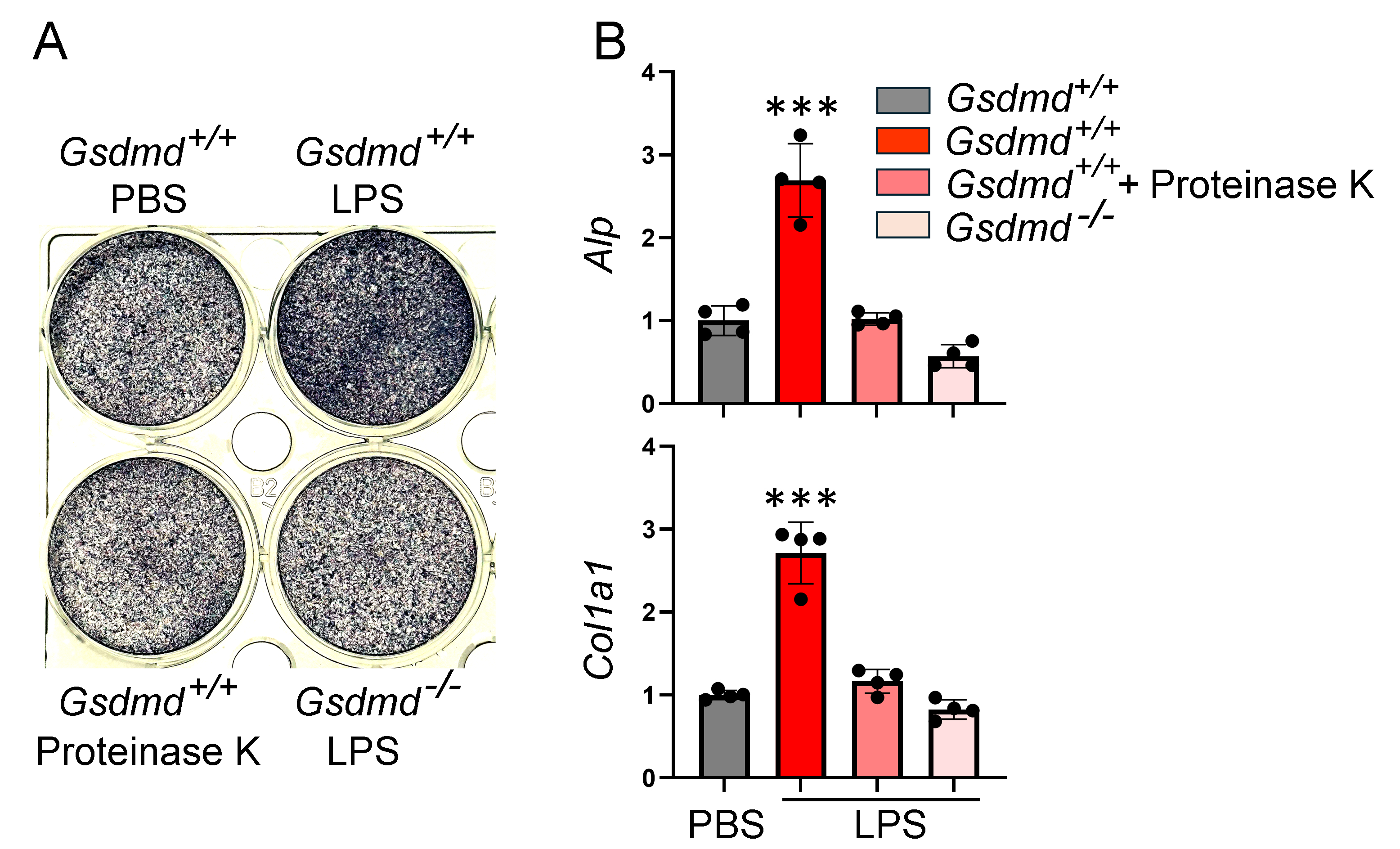

**Supplementary Fig. 9. Bone marrow supernatants from LPS-treated mice promote osteogenesis *in vitro*, a response that is GSDMD-dependent.** Bone marrow cells and **p**eriosteal cells from WT mice were expanded for 7-10 days and cultured in osteogenic medium (50 μg/ml of ascorbic acid and 2 mM of glycerol 2-phosphate). Cultures were supplemented with bone marrow supernatants collected on day 3 from *Gsdmd^+/+^* and *Gsdmd^-/-^* mice treated with PBS or LPS on days 0. In some conditions, WT supernatants were treated with proteinase K prior to use. **A**) Alkaline phosphatase activity. **B**) qPCR analysis. ***p<0.001. One way ANOVA.

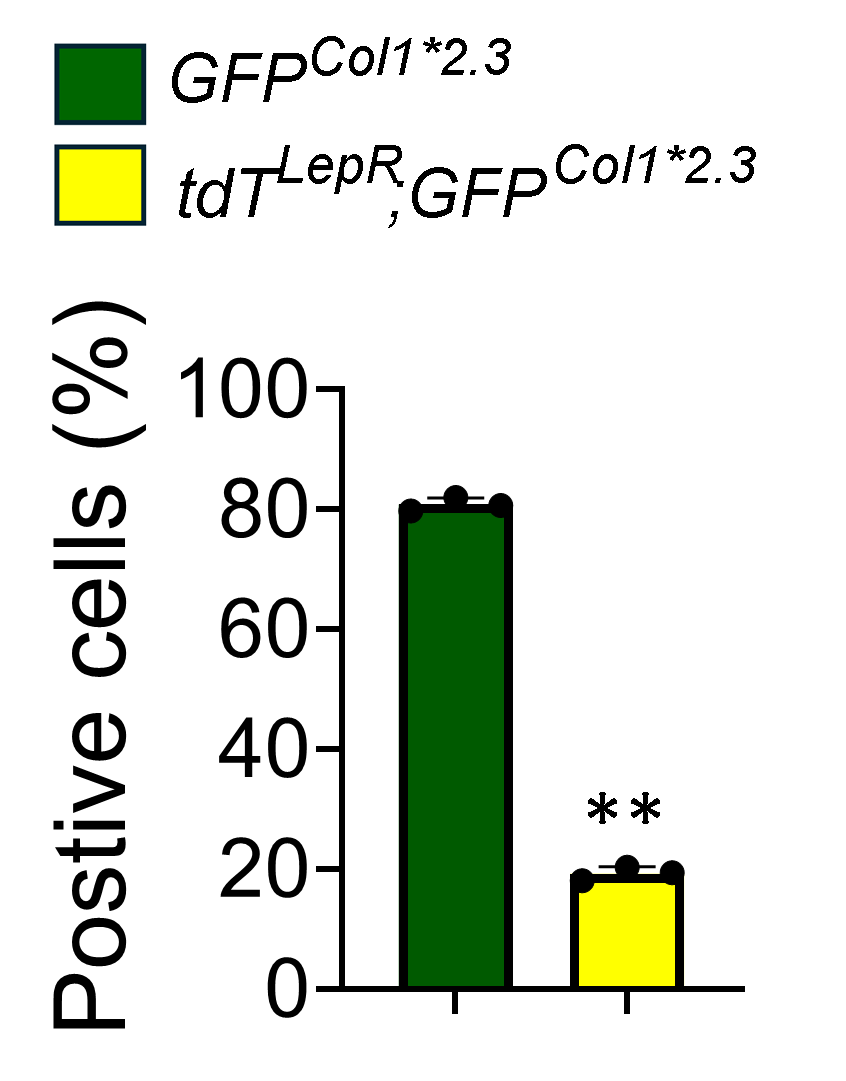

**Supplementary Fig. 10. LepR+ cells** **differentiate into OBs in response to LPS treatment**. Three months old *tdT^LepR^*;*GFP^Col1*2.3^* mice were i.p. injected with 1 mg/kg LPS (n=3) on day 0, 4, and 8 and analyzed on day 10 as described in Materials and Methods. tdT^LepR^;GFP^Col1*2.3^ cells and GFP^Col1*2.3^ cells in periosteum were analyzed.
